# Inferring Functional Neural Connectivity with Deep Residual Convolutional Networks

**DOI:** 10.1101/141010

**Authors:** Timothy W. Dunn, Peter K. Koo

## Abstract

Measuring synaptic connectivity in large neuronal populations remains a major goal of modern neuroscience. While this connectivity is traditionally revealed by anatomical methods such as electron microscopy, an efficient alternative is to computationally infer functional connectivity from recordings of neural activity. However, these statistical techniques still require further refinement before they can be reliably applied to real data. Here, we report significant improvements to a deep learning method for functional connectomics, as assayed on synthetic ChaLearn Connectomics data. The method, which integrates recent advances in convolutional neural network architecture and model-free partial correlation coefficients, outperforms published methods on competition data and can achieve over 90% precision at 1% recall on validation datasets. This suggests that future application of the model to *in vivo* whole-brain imaging data in larval zebrafish could reliably recover on the order of 10^6^ synaptic connections with a 10% false discovery rate. The model also generalizes to networks with different underlying connection probabilities and should scale well when parallelized across multiple GPUs. The method offers real potential as a statistical complement to existing experiments and circuit hypotheses in neuroscience.

## 1. Introduction

Precise brain wiring diagrams can be used to validate and generate hypotheses of neural circuit function. In recent years, considerable resources have been devoted to ultrastructural reconstructions of synaptic connectivity from serial section electron microscopy (EM) data (Briggman et al., 2011; Chalfie et al., 1985; Hildebrand et al., 2017). But these anatomical methods, while improving, remain slow, costly, and laborious. An emerging alternative to EM reconstruction is statistical inference of the synaptic connectivity latent in observed neural activity patterns. Various statistical methods for inferring connectivity have been published, ranging from model-free approaches, such as Granger causality (Cadotte et al., 2008; Garofalo et al., 2009), transfer entropy (Stetter et al., 2012), and partial correlation coefficients (Sutera et al., 2014), to model-based Bayesian approaches that seek to invert generative models of neural activity (Mishchenko et al., 2011; Soudry et al., 2015).

One critical issue with connectivity inference is that two unconnected neurons with common inputs will have correlated activity that is easily interpreted as causal if the common input is unobserved. A recent method proposes a generalized linear model-based Bayesian method to combat the common input problem when the analysis is coupled with a “shotgun” imaging experiment that hypothetically measures activity in all neurons serially (Soudry et al., 2015). The method is accurate and efficient when prior information about connection sparsity is known and when activity in the neural population of interest behaves according to the dynamics assumed by the generative model. In practice, however, these assumptions can be problematic, especially as the true connection probabilities in real brain networks are hard to measure.

The common input problem ceases to be an issue in neural systems with complete observability. Such universal access has been granted by recent technological advances enabling cellular resolution fluorescence imaging of neural activity on large scales (Chen et al., 2013). In particular, the activity of nearly all neurons in the larval zebrafish brain can be imaged at speeds up to 12 Hz (Ahrens et al., 2013; Dunn et al., 2016; Tomer et al., 2015). While these experiments skirt the common input problem, the acquired data presents a unique set of challenges for connectivity inference, as action potentials are obscured by the noise and slow dynamics of fluorescent indicators. Nevertheless, there have been attempts to extract meaningful connectivities from these recordings. Recently, several methods were published together as part of the Kaggle ChaLearn Connectomics competition (Orlandi et al., 2014). Here, we report an improved deep learning method that outperforms the competition leader and, importantly, provides a viable false discovery rate (FDR) at a true positive rate (TPR) poised to reveal millions of synaptic connections in real brain volumes.

For clarity, throughout this report we will refer to neurons in the synthetic networks that emit measurable fluorescence signals as “cells” and refer to those synthetic networks as “graphs.” The neural networks used for prediction and classification will be referred to as “networks” or “models.”

Code for the following models and analyses is available at https://github.com/spoonsso/TFconnect.

## 2. Related Work

Convolutional neural networks (CNNs) are considerably powerful for computer vision, automatically extracting spatially invariant, hierarchical features and performing classification within an end-to-end learning framework (Krizhevsky et al., 2012). Outside of the computer vision domain, CNNs were recently adapted to analyze fluorescence time series from cells in connected graphs, with spatiotemporal filters operating across cells over time (Fig. 1a) (Romaszko, 2015). These networks learned to classify pairwise binary connectivity when trained on activity generated from synthetic graphs of 1000 cells, using binary labels from the associated *in silico* connectivity matrix.

**Fig 1:**
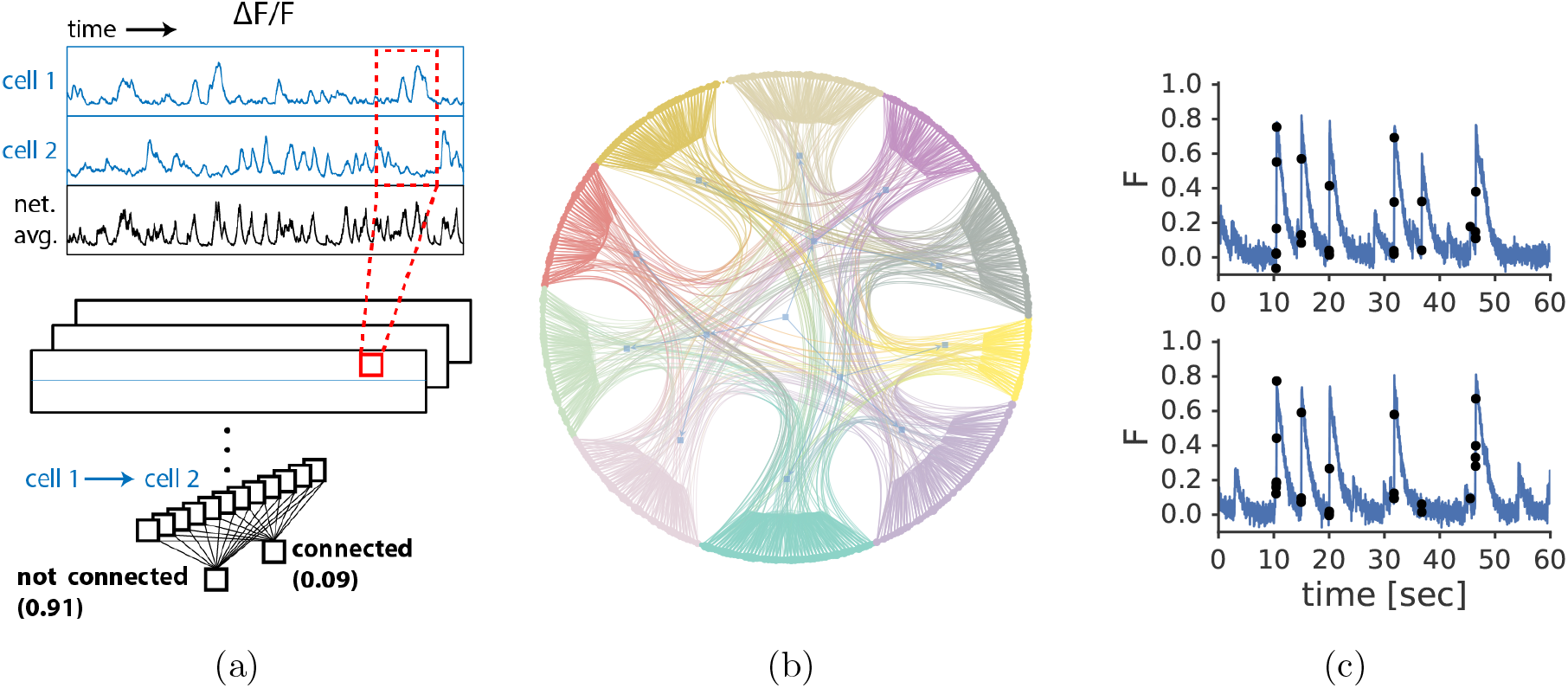
Overview of the problem and method. (a) We train a deep convolutional neural network to predict the underlying connectivity of cells. (b) The graphs consists of 1000 cells separated into 10 sub-graphs (different colors) with higher clustering coefficients. For clarity, only 400 cells are shown. (c) The goal of the method is to predict the presence or absence of a connection between each cell pair in the graph based only on the observed activity-generated fluorescence in each cell. The method operates on signals subsampled at time points where the overall activity across cells in the graph is high (black dots).

Over the past three years, a host of improvements to CNN architectures have been published. These improvements include dropout (Srivastava et al., 2014), where a random subset of connection weights are temporarily withheld over a mini-batch during training in order to combat overfitting, batch normalization (Ioffe and Szegedy, 2015), where the pre-activated output of each layer is re-normalized before being passed to the next layer in order to improve training efficiency, and residual blocks (He et al., 2015), which drastically increase the effective depth and therefore the expressivity of the CNN. Furthermore, parametric rectified linear unit (PReLU) activations allow the network to learn a suitable leakiness in its non-linear rectification activation, and intelligent weight initialization has been shown to increase training efficiency (He et al., 2016).

In parallel, several alternative methods for inferring functional connectivity have been published. One method, using a partial correlation coefficient metric estimated from the inverse covariance matrix (Sutera et al., 2014), set the state-of-the-art benchmark at the inaugural Kaggle ChaLearn Connectomics competition, edging out baseline summary statistics like correlation coefficients and entropy-based causality estimations (Orlandi et al., 2014).

We find that by incorporating these advances in deep network design, and by combining partial correlation measurements with fluorescence traces, we can dramatically increase the performance of a residual CNN (RCNN) model for connectomics analysis. While we focus primarily on the Kaggle ChaLearn competition datasets, as they enable direct comparisons across several methods, we will discuss our work in the context of formal Bayesian approaches that attempt to infer underlying connectivity matrices via inversion of generative models. Compared to these approaches, we suggest that our RCNN method is more robust to variation in graph connection probability and will be much more scalable.

## 3. Synthetic Graph Architecture

The Kaggle ChaLearn Connectomics data are generated from a realistic model of neural dynamics (leaky integrate-and-fire), calcium binding, and fluorescence (Stetter et al., 2012; Orlandi et al., 2014). Graph model parameters were tuned so that cells were spontaneously active, including periods of pan-graph bursting. These parameters and associated dynamics were chosen by the competition organizers to closely resemble real networks of primary cultured neurons. In the future, we aim to train on synthetic data generated from graphs whose architectures and dynamics better resemble that of brains in *vivo*. That being said, the existing competition dataset does include realistic features that are likely shared by the brains of real animals.

In our study, we used signals from cells in 8 different graphs for the purposes of training, validation, testing, and analysis. Six of the graphs (which we call “normal”) had the same average architecture, with 3 used for training sets, 1 for validation, and 2 for testing. Each of these graphs contained 1000 cells with an overall average connection probability *ρ* = 0.014 (Fig. 1b). The graph comprised 10 subgraphs whose cells had higher within group connection probabilities (*ρ_int_* = 0.012) than between group connection probabilities (*ρ_ext_* = 0.002). The weights of connections, which were all positive/excitatory, were the same within a given subgraph, and the weights in each subgraph were ultimately optimized to produce a desired universal bursting frequency of 0.1 Hz.

## 4. Residual Convolutional Neural Network Architecture

### Preprocessing of fluorescence signals

Before training, signals were subjected to three rounds of preprocessing in order to accentuate information content and reduce overall data size. First, signal noise arising from simulated light scattering effects was removed via spatial deconvolution (physical locations of each cell were provided as part of the dataset). Second, each signal was downsampled by thresholding a high-pass filtered representation of the total graph activity (Fig. 1c). Third, each downsampled signal was z-scored by subtracting the downsampled signal mean and dividing by the downsampled signal standard deviation.

### Structuring training input

Following the general scheme of (Romaszko, 2015), these processed signals were then assembled into chunks of 3x330 continuous samples, beginning at random starting points in the processed signals. The first two rows of this data structure contained processed signals from a pair of cells, with the associated label referencing the binary directed connection between the cell in row 1 and the cell in row 2. The third row contained the corresponding average fluorescence across the entire graph for the time interval included in rows 1 and 2 (Fig. 1a).

The final training set included approximately 1.1 million paired examples in this format, with equal representation of positive and negative examples (i.e. the presence or absence of a directed connection, respectively). Note that 550,000 far exceeds the total number of positive examples contained in 3 graphs (~ 42,000), but a rich, unique training set can be assembled because the starting position for each 330-sample training chunk is random for each example. Of the ~ 1.1 million examples in the training set, 75% were used for training epochs and the other 25% were used as cross-validation.

### Residual convolutional neural network architecture

The RCNN begins with input [3 × 330 × 1], contains 1 block of (1 + 2) convolutional + residual convolutional layers (each [2 × 326 × 32] with filters [2x5x1]), another 1 block of (1 + 2) convolutional + residual convolutional layers (each [1 × 322 × 64] with filters [2 × 5 × 32]) followed by [1 × 10] max pooling, another full convolutional layer (size [1 × 32 × 128] with filters [1 × 1 × 64]), 1 block of (1 + 2) dense + residual dense layers (each with 256 units), and a dense readout layer (2 units) indicating the final softmax classification (Fig. 2a). Each layer implemented PReLU activations.

**Fig 2:**
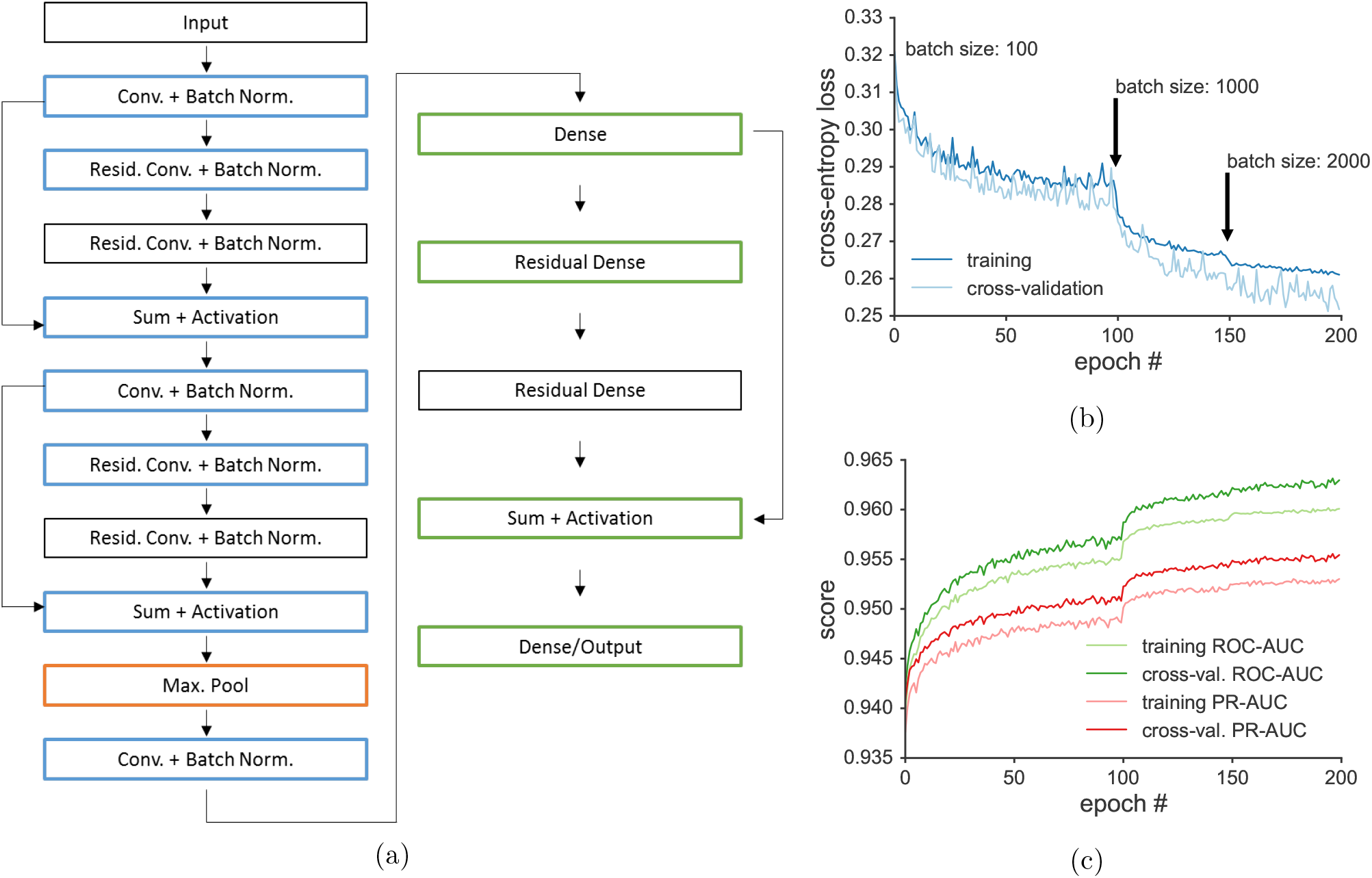
Details of the RCNN and training. (a) Summary of layer architecture. (b) Training time-course for a single model. Where indicated, the batch size was annealed to promote learning. (c) Training time-course but for area under the receiver operator characteristic and precision recall curve metrics (ROC-AUC and PR-AUC, respectively). Note that the typical number of training epochs was 200 for each batch period (this example shows only 100/50/50).

### Training

The model was trained using a cross-entropy loss function based on proper binary classification of each cell pair’s connectivity (Fig. 1a, 1c). For each individual model, we started with a batch size of 100 and performed a maximum of 200 epochs with an automatic early stopping criterion: if the best cross-validation loss did not improve over a span of 20 epochs, training halted. We then annealed the batch size, progressing to 1000 examples and then 2000 examples per batch. These extra steps consistently resulted in large, sudden reductions in training loss (Fig. 2b) and gains in performance metrics (Fig. 2c). Gradient descent was performed using Adam optimization using recommended default parameters (Kingma and Ba, 2014). We employ *l*2 weight decay with a regularization parameter set to 10^−6^.

The RCNN was implemented in Python using tfomics (https://github.com/spoonsso/tfomics), a high-level API for TensorFlow (v0.12.1). On a single NVIDIA Titan X Pascal graphics card, 1 epoch during the 100 batch phase took 434 s (50 ms / batch) including a forward pass for crossvalidation, during the 1000 batch phase took 241 s (280 ms / batch), and during the 2000 batch phase took 245 s (569 ms / batch).

During training, we also introduced dropout at each of the layers in the RCNN. For the convolutional layers, the dropout probability was set to 0.2, and for the dense layers, the dropout probability was set to 0.5. At each layer, we included batch normalization of parameters. At the beginning of training, weights were initialized following the distribution outlined in (He et al., 2016). Taken together, these measures help to reduce overfitting and increase training efficiency.

### Testing

Similar to (Romaszko, 2015), we completed 14 forward passes through the trained network for each cell pair, tiling the entire length of the signals in 330 sample patches. The predicted scores were then averaged across each of these 14 passes to arrive at a final prediction for connectivity between the two cells. Forward passes took 1.3 ms per cell pair, for a total of ~21.7 minutes to evaluate the entire connectivity matrix. We then ensemble averaged the predictions for each cell pair from 5 independent models trained separately, with differences in final model parameters resulting from the random initialization of weights and the random assembly of network input from training data.

## 5. Results

### The RCNN sets competition benchmark

When we evaluated the RCNN performance on the Kaggle ChaLearn Connectomics competition test datasets, it outperformed all other entries in the area under the receiver operator characteristic (ROC) curve (Table 1). Examination of full ROC curves on the “normal” validation graph (*ρ* = 0.014) revealed that while the model ensemble outperformed any individual model in total area under the curve (AUC), some individual models performed better than the ensemble locally within small ranges along the curve (Fig. 3a). The RCNN model also outperformed the winning algorithm, which was based on partial correlation coefficients (PCs) between cells across the dataset, on a denser validation graph with a higher average connection probability *ρ* = 0.022. However, the RCNN model struggled on a sparser validation graph with a lower average connection probability *ρ* = 0.007 (Table 1).

**Fig 3:**
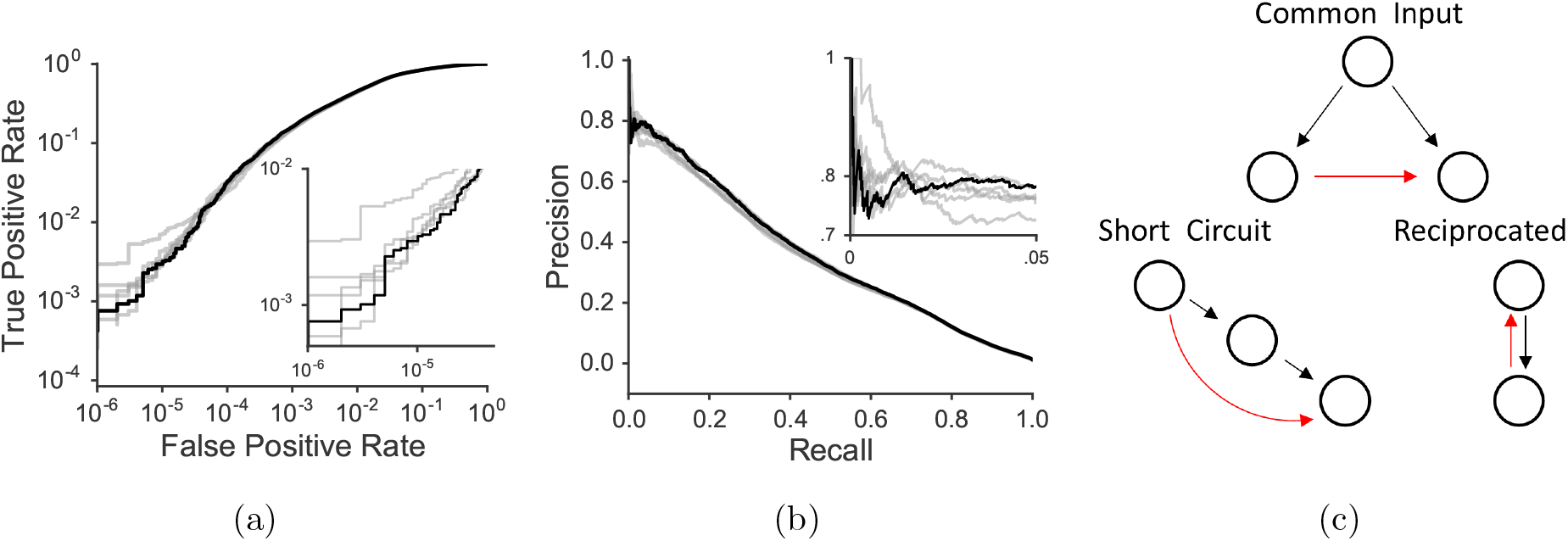
Performance of the RCNN. (a) ROC curves on a validation dataset for 5 individual models (gray lines) and a final ensemble-averaged model (black line). (b) PR curves for the same models. (c) Schematics of three problematic connectivity motifs with which the model struggles. Black arrows represent true connections and red arrows denote inferred false positives.

**Table 1.** Comparison of algorithms on predictions for five networks (graphs) from the Chalearn connectomics challenge. Area under the ROC and PR curves is compared. PR curves for Test Networks 1 and 2 are unavailable as the data are managed by Kaggle. RCNN + PC sets benchmarks in all networks and metrics.

| Algorithm | Valid Network |  | Test Network 1 |  | Test Network 2 |  | Dense Network |  | Sparse Network |  |
| --- | --- | --- | --- | --- | --- | --- | --- | --- | --- | --- |
|  | ROC | PR | ROC |  | ROC |  | ROC | PR | ROC | PR |
| corr. coef. | 0.681 | 0.026 | 0.664 |  | 0.689 |  | 0.595 | 0.030 | 0.691 | 0.023 |
| PC | 0.939 | 0.388 | 0.942 |  | 0.941 |  | 0.942 | 0.413 | 0.955 | 0.334 |
| CNN | n.d. | n.d. | 0.940 |  | 0.941 |  | n.d. | n.d. | n.d. | n.d. |
| RCNN | 0.943 | 0.359 | 0.943 |  | 0.944 |  | 0.952 | 0.383 | 0.910 | 0.222 |
| RCNN + PC | **0.953** | **0.449** | **0.951** |  | **0.952** |  | **0.954** | **0.456** | **0.955** | **0.372** |

### Evaluating the model’s realistic potential

The ultimate goal of our method is to infer, with an acceptable degree of precision, real synaptic connectivity from real neural imaging data. Examination of the model’s precision recall (PR) curve on the validation dataset suggests that RCNNs could achieve a stable level of ~80% precision (~20% FDR) at up to 5% recall (Fig. 3b). For a real zebrafish brain imaging dataset with 10^5^ cells and 10^10^ possible connections with ~ 2% connectivity (Stobb et al., 2012), even 5% recall would report 10^7^ connections, with 1 in 5 being false. At an FDR of 10%, the RCNN recalls 0.07% of the connections, or ~ 10^5^ connections in a typical zebrafish brain.

To gain a better understanding of where the RCNN model fails, we quantified the frequency of three potentially problematic connectivity motifs. Because the RCNN operates only on pairwise input, the model is particularly susceptible to misclassifying correlated activity from common inputs as a directed causal relationship (Fig. 3c, top). Similarly, and enhanced by the slow indicator dynamics and acquisition rate, the RCNN might also mistake a polysynaptic connection for a directed monosynaptic connection between the first and last cell in a chain (Fig. 3c, bottom left, “short circuit”). Finally, without access to millsecond spike times, it should be difficult for a model operating on slow and noisy fluorescence signals to infer the direction of causality in correlated activity (Fig. 3c, bottom right). Indeed, 47% of all possible common input relationships, 49% of all possible “short circuit” motifs, and 71% of all possible “reciprocated” connectivities were classified as false positives (Table 2). The model especially struggled when two or more of these motifs were combined, with 98% of all possible false positives shared by all 3 motifs being classified erroneously.

**Table 2.**
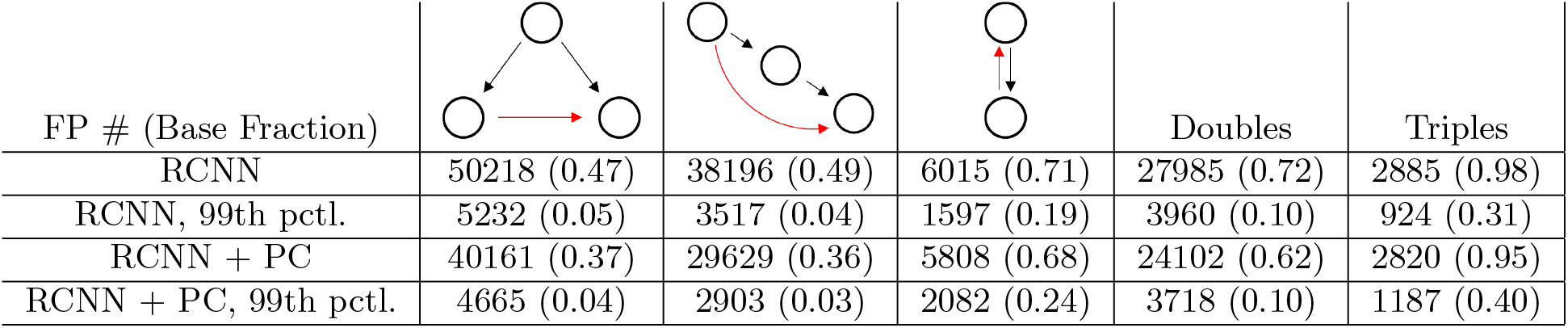
Analysis of predicted false positives (FPs) for the original RCNN model and, the improved RCNN model incorporating partial correlation coefficients. For each of the FP types (icons left to right: common input, short circuit / polysynaptic, reciprocated), and, combinations thereof (overlapping double and triple motifs), the total number of false positives and fraction of the total possible base frequency of each type are indicated. FPs are evaluated using either forced choice or at a 99^th^ percentile prediction score threshold.

| FP # (Base Fraction) |  |  |  | Doubles | Triples |
| --- | --- | --- | --- | --- | --- |
| RCNN | 50218 (0.47) | 38196 (0.49) | 6015 (0.71) | 27985 (0.72) | 2885 (0.98) |
| RCNN, 99th pctl. | 5232 (0.05) | 3517 (0.04) | 1597 (0.19) | 3960 (0.10) | 924 (0.31) |
| RCNN + PC | 40161 (0.37) | 29629 (0.36) | 5808 (0.68) | 24102 (0.62) | 2820 (0.95) |
| RCNN + PC, 99th pctl. | 4665 (0.04) | 2903 (0.03) | 2082 (0.24) | 3718 (0.10) | 1187 (0.40) |

### Improving the model

We reasoned that many of these errors could be eliminated if the RCNN could make multivariate evaluations that considered the activity across all cells in the graph rather than just pairwise correlations. Partial correlation coefficients (PCs) are multivariate summaries of causality and led the overall competition leaderboard (Sutera et al., 2014). Reasoning that the strengths of PC-based classification and our RCNN model might be complementary, we incorporated the PC for each cell pair into our input structure as a fourth row of data during evaluation and training. Other than adjusting the size of input layer and the first residual block to accommodate the PC addition, this RCNN + PC model used the same gross network topology.

The final RCNN + PC model, which was an ensemble average of 8 models, including 1 model trained on data with an unequal representation of positive and negative examples (10% and 90%, respectively), significantly improved the ROC curves on the validation dataset (Fig. 4a) and the ROC-AUC and PR-AUC metrics on all validation and test datasets (Table 1).

**Fig 4:**
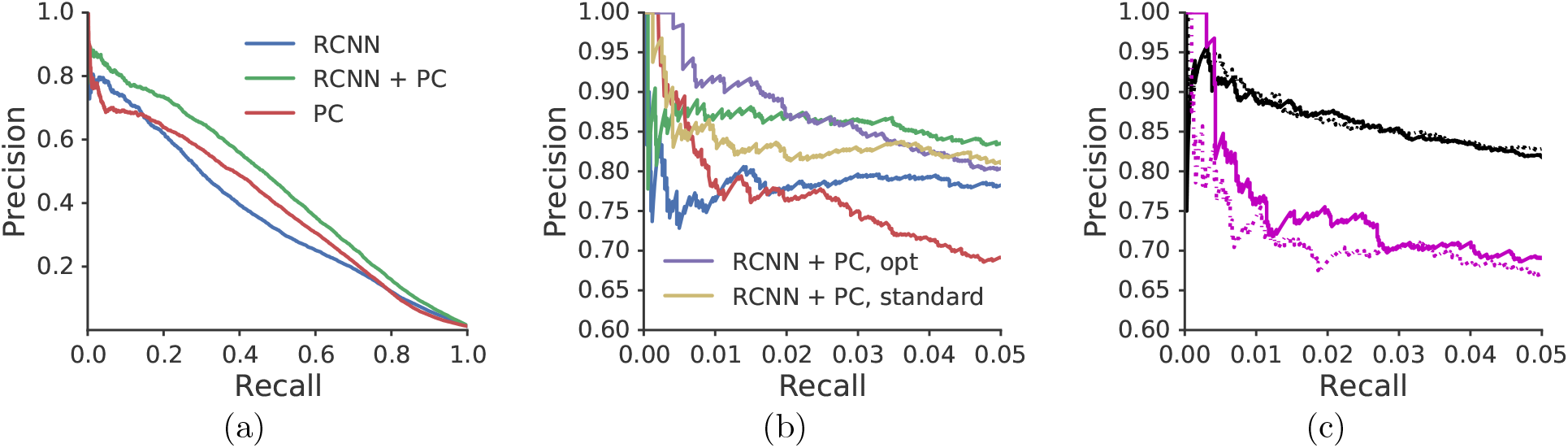
Improvement of predictions in precise regimes. (a) Full PR curves on a validation dataset for the original RCNN, a new RCNN that incorporates multivariate partial correlation (RCNN + PC), and PC alone (b). Zoomed-in PR curves for the same models, plus a model with pre-ensemble weight optimization (opt) and an ensemble average excluding a model with low representation of positive examples (standard). (c) Full ensemble (solid) and “standard” ensemble (dashed) PR curves for additional dense (black) and sparse (magenta) connectivity validation data.

Comparing the precise regions of the PR curves across algorithms reveals that the RCNN + PC model provides an increase in the precision plateau at the start of the curve, maintaining over 85% precision past 3% recall, a significant improvement over the original RCNN model and the PC algorithm alone (Fig. 4b). Note that at up to 0.5% recall, the PC algorithm shows enhanced precision over the RCNN + PC model, but this may be due to the incorporation of predictions from the model trained on 10%/90% positive/negative examples; without this model, the RCNN + PC ensemble shows enhanced precision at low recall but does not sustain this precision for long. We were also able to enhance the early precision (from 0 to 2% recall) of the full RCNN + PC ensemble using a Gaussian process search for each model’s weight.

These improvements in PR reflect, at least in part, reductions in the misclassification of some problematic connectivity motifs (Table 2). Overall common input false positives (FPs) were reduced by 20%, “short circuit” FPs by 22%, “reciprocated” FPs by 3%, doublet FPs by 14%, and triplet FPs by 2%. For the top 1% of the predictions, however, these gains were somewhat muted. In this regime, “reciprocated” FPs actually increased in the RCNN + PC, perhaps indicating the dominance of the PC representation, which is agnostic to edge direction, in this regime.

## 6. Model Generalization

Effective generalization is a major consideration for future applications of these models to real data. Any model trained on data generated from a limited set of synthetic graphs will likely need to operate on signals arising from a graph with a different underlying parameter distribution, even if the models can eventually be trained on ground truth labeled data. It is thus important that the predictive power of the model generalize to validation graphs with different architectures that were not included in the training set. We thus tested the trained RCNN + PC ensemble on the dense and sparse validation datasets. For the sparse graph, the model was reduced in effectiveness, achieving on 75% precision at 1% recall, although there continued to be regimes where the model achieved high levels of precision. For the dense graph, the model performed better, achieving almost 95% precision at 1% recall (Fig. 4c). Note that the connection probability of the dense graph is close to the anticipated average connection density of the zebrafish brain (Stobb et al., 2012).

Successful generalization also requires that a desirable prediction threshold deduced from analysis on validation data be effective when applied to real data, where PR curves cannot be plotted in search of a viable threshold. While the distribution tails of prediction scores depend on connection sparsity, (Fig. 5a), we find that recall and precision on the dense and sparse validation data can be recovered by choosing thresholds matched to prediction score percentiles in the “normal” validation dataset (Figs. 5b, 5c).

**Fig 5:**
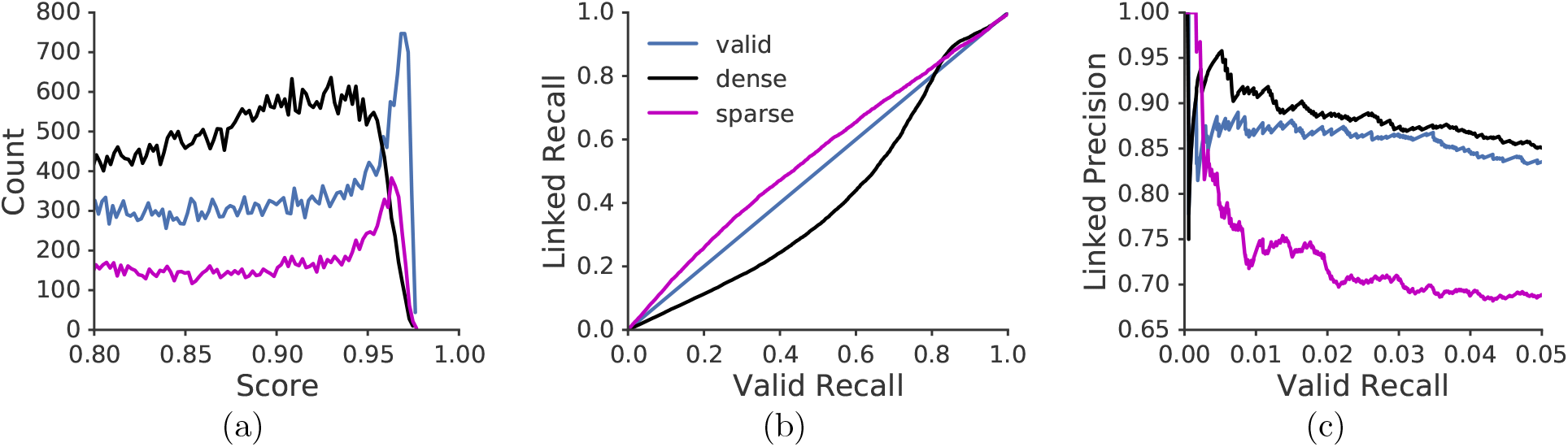
Generalization of the model to graphs with different structure. (a) Histograms of score distribution tails for each graph. (b) Model recall for densely and sparsely connected validation graphs as a function of recall on a validation graph whose density matched the training data. (c) Precision for each graph as a function of validation recall.

## 7. Scalability

The evaluation times for the trained RCNN + PC model have time complexity *O*(*n*^2^), as the network independently operates on all cell pairs in the dataset. With a current evaluation time for 1.3 seconds for a single cell pair, we anticipate that applying the method on a full fish dataset of 10^5^ cells will require 3611 GPU hours. This evaluation process is easily parallelized, however, as independent GPUs can load the final model parameters and evaluate batches of input independently. On a 128 GPU cluster, this process would take 28.2 hours for a single model running perfectly parallel. While this evaluation would need to be run multiple times, once for each model in the ensemble, we note that we found a single model that outperformed the ensemble on the competition test ROC-AUC metric. Future work should determine the utility of single-model solutions to reduce compute time. We also note that these times reflect evaluation on an entire brain. Analyses of local connectivity in smaller brain nuclei would be considerably faster.

For larger datasets, PCs can also be used to pre-screen for connected candidates, reducing the overall number of cell pairs that need to be evaluated by the RCNN. For instance, 100% of the predicted true positives at 3% recall on the”normal” validation dataset can be recovered by preselecting for cell pairs that are in the 99^*th*^ PC percentile, reducing the expected evaluation time to ~36 hours for 10^5^ cells on a single GPU, or ~17 minutes on a 128 GPU cluster. We extrapolate that the PC calculation for an hour-long, 5 Hz recording of 10^5^ cells would take ~14 hours on a single 6-core machine.

## 8. Discussion

### Improvements to the RCNN architecture

Future work should explore a larger space of layer topology and hyperparameters in order to test the limits of classification efficacy. Deeper models may improve the overall performance and efficiency of the training process. The input data structure may also benefit from an expansion in the total number of samples passed to the network during evaluation and training. Longer traces, which contain more information about the temporal relationships between cells, may also improve the overall performance of the model.

The model should, in principle, be at least as good as the PC metrics passed to the network alongside the fluorescence signals, but in early regimes of the PR curve, the PC algorithm alone appears to be more accurate. In order to provide more salience to the PC score, it may help to pass PC coefficients through the model separated from the learned convolutional filters, as PC coefficients have no spatiotemporal correspondence to cell activity. Future model designs should also incorporate better multivariate representations of cell activity throughout the graph. If the model were able to operate on more than just one cell pair at a time, it could in principle learn representations of higher order correlational structure.

In order to combat problematic connectivity motifs, it may help to over-represent them in the training dataset. In the final RCNN + PC ensemble, we included a model that was trained on 10%/90% positive/negative examples rather than 50%/50% with the hope that including more FP examples might help the network distinguish FPs from true positives more easily. While we have not explored this configuration exhaustively, that model appeared to contribute greater precision in the 1-5% recall range at the expense of precision in the 0-1% recall range. Future work should explore the extent to which models trained on different proportions of positive and negative examples can complement each other and enhance overall classification accuracy.

### Generalization to non-synthetic neural data

While ground truth labeled data may become available as EM and all-optical circuit mapping techniques progress, the success of the current method will rely on synthetic training data reflecting realistic models of neural dynamics. Studies of neural network generalization from synthetic to real data (Jaderberg et al., 2014; Le et al., 2017) suggest that training on synthetic data can result in powerful generalization, given that the networks are trained on a large enough space of synthetic examples. Going forward, it will be critical to simulate neural activity and concomitant fluorescence signals that better reflect the activity observed *in vivo*. We also note that generalization of the RCNN model will depend on what it has learned to detect in the fluorescence traces. It is unclear how much the learned convolution filters depend on the specifics of the observed neural dynamics and thus the underlying parameters of the generative model of simulated activity. It’s possible that the RCNN model is learning a model-free causality statistic that will generalize well even to neural dynamics that differ substantially from the training data. Future saliency analyses and probes of RCNN network representation will thus speak to the method’s overall potential for generalization.

Ultimately, the vision is to train the RCNN on synthetic data closely matching the dynamics observed spontaneously in larval zebrafish (Dunn et al., 2016). Conservative predictions, thresholded at a percentile reflecting a target confidence in precision or recall, can be cross-validated against known larval zebrafish brain anatomy and eventually validated via optical or ultrastructural circuit mapping methods, such as channelrhodopsin/GCaMP (Packer et al., 2015), PA-GFP (Dunn et al., 2016), and EM (Hildebrand et al., 2017). The idea is not that the current method will be able to uncover the complete connectome of an animal but rather that it will be able to lend statistical support to circuit-level hypotheses of neural structure and function.

One advantage of this method over Bayesian techniques for inferring functional connectivity is that it works directly on calcium imaging data, avoiding slow and potentially error-prone inference of spike timing from fluorescence signals. Furthermore, full Bayesian inference of adjacency matrices is slow, and faster approximations typically make strong assumptions about underlying connection sparsity (Soudry et al., 2015), a property that is not straightforward to estimate in real systems. In contrast, our method is relatively robust to changes in connection probability.

Finally, we note that the precision of the method is agnostic to the types of underlying FPs. While it will be important in the future to better combat problems related to common inputs in order to improve the overall accuracy of the method, the RCNN + PC model is applicable in its current form as long as one accepts the uncertainty of a target FDR. As in genomics analyses, a target FDR of 5 to 10% should be reasonable assuming the researcher is aware of its implications.

### Summary

We present here a new deep learning method for inferring functional connectivity from recordings of fluorescence from calcium-dependent activity indicators. The method sets an important state-of-the-art benchmark, as evaluated on the Kaggle ChaLearn Connectomics competition, and shows immediate promise for applications to real data. Going forward, a comprehensive understanding of the constraints and pitfalls of the method will be critical for establishing reliable, conservative predictions of functional connectivity.

## Acknowledgments

This work was supported by an NVIDIA hardware grant (Titan X Pascal).

